# Adaptive sampling-based structural prediction reveals opening of a GABA_*A*_ receptor through the *αβ* interface

**DOI:** 10.1101/2024.05.03.592340

**Authors:** Nandan Haloi, Rebecca J. Howard, Erik Lindahl

## Abstract

GABA_***A***_ receptors are ligand-gated ion channels in the central nervous system with largely inhibitory function. Despite being a target for drugs including general anesthetics and benzodiazepines, experimental structures have yet to capture an open state of canonical ***α***1***β***2***γ***2 GABA_***A***_ receptors. Here, we use a goal-oriented adaptive sampling strategy in molecular dynamics simulations followed by Markov state modeling to capture an energetically stable putative open state of the receptor. The model conducts chloride ions with comparable conductance as in electrophysiology measurements. The channel desensitizes by narrowing at both the cytoplasmic (**−2**^**′**^) and central (**9**^**′**^) gates, a motion primarily mediated by transmembrane ***αβ*** subunit interface. Consistent with previous experiments, targeted substitutions disrupting interactions at this interface slowed the open-to-desensitized transition rate. This work demonstrates the capacity of advanced simulation techniques to investigate a computationally and experimentally plausible functionally critical of a complex membrane protein, yet to be resolved by experimental methods.

## Introduction

*γ*-Aminobutyric acid type A (GABA_*A*_) receptors are pentameric ligand-gated ion channels that mediate fast inhibitory synaptic transmission in the vertebrate central nervous system. The neurotransmitter GABA, upon release at the synaptic cleft, binds to the extracellular domain (ECD) of these receptors in the resting/closed state. This triggers an allosteric signal to the transmembrane domain (TMD) to transiently open the pore for the selective flow of chloride ions across the plasma membrane, before the channel enters a desensitized state refractory to activation upon sustained GABA binding (Fig. 1A). This process is proposed to be regulated by a “dual gate mechanism”: 1) the upper half of the pore, 9^′^ gate, is shut in the resting state, 2) upon activation the contraction and the rotation of ECD (‘unblooming’ and ‘twisting’) widens the 9^′^ gate and opens the pore, and then 3) a gate located at the intracellular end of the pore, −2^′^ gate, closes during desensitization [1] (Fig. 1B).

**Fig. 1.**
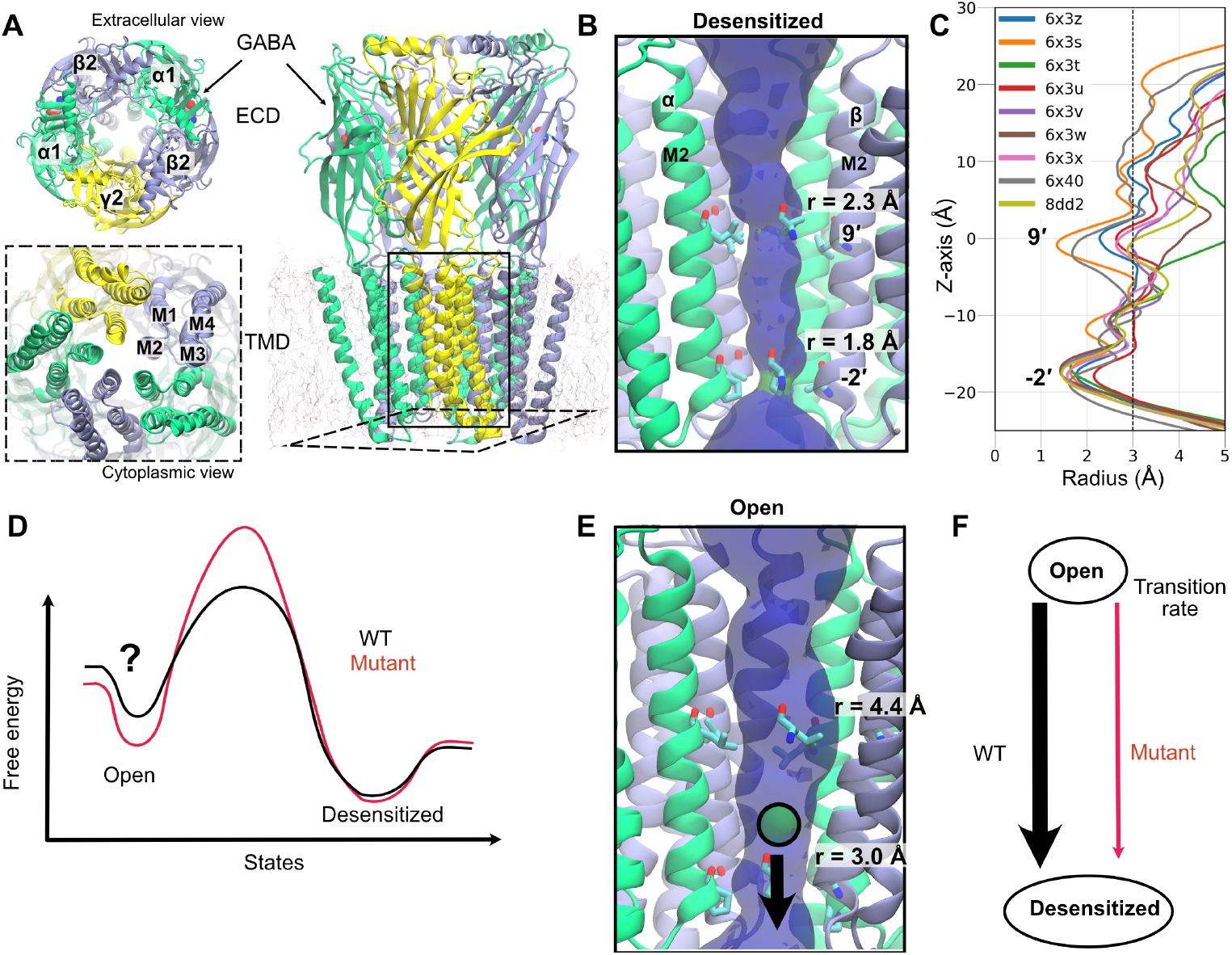
Overview of structural, energetic and kinetic aspects GABA_*A*_ receptor functions explored in this work. A) Architecture of a representative GABA_*A*_ receptor, colored by subunit (*α* in green, *β* in blue, *γ* in yellow), viewed from the extracellular (*left, top*), cytoplasm (*left, bottom*) and membrane plane (*right*). Horizontal slice through TMD shown from the cytoplasmic perspective (*left, bottom*), highlighting the 4 helices where M2 faces the pore. Two GABA-bound at the orthosteric site between the *α*-*β* submit is represented in van der Waals spheres. B) The ion permeation pathway (determined using the program HOLE [22]) of the GABA_*A*_ receptor at the desensitized state (PDB ID: 6×3z), depicted with blue and green colors, the latter representing a narrowness of the pore where there is room for a single water. For clarity, the *γ* subunit was removed. The residues located at the two gates are highlighted; 9^*′*^ is formed by a leucine residue at each subunit and −2^*′*^ is formed by a proline at the *α* and *γ* and by an alanine at the *β* subunit. Also, the radius at these two gates is shown. C) Pore radius profile of *α*1*β*2*γ*2 GABA_*A*_ receptor structures, either in the closed or desensitized state, bound to different ligands [6]. 6×3z contains GABA, 6×3s bicuculline, 6×3t GABA+propofol, 6×3u GABA+propofol, 6×3v GABA+etomidate, 6×3w GABA+phenobarbital, 6×3x GABA+diazepam, 6×40 GABA+picrotoxin, 8dd2 GABA+zolpidem. The vertical dashed line represents the radius of a hydrated chloride ion. The radius profile of *α*1*β*3*γ*2 GABA_*A*_ receptor structures can be found in Fig. S1 [10, 11]. D) A conceptually simplified free energy profile, highlighting the fact that the open state is relatively less stable than the desensitized state. The red line corresponds to a mutant that was characterized in this study that can prolong the desensitization process by increasing the energy barrier between these two states. E) The pore profile of the open state discovered in this study, high-lighting the radius at the two gates. A chloride ion (in green) is depicted that can readily permeate through this pore, as shown in our results. F) A schematic two-state kinetic model of the WT and a mutant system, that was calculated in this study and compared with experimental results.

Nineteen human GABA_*A*_ subunits have been has been identified so far that includes *α*1–6, *β*1–3, *γ*1–3, *δ, ϵ, θ, π*, and *ρ*1–3 [2–4]. In the brain, most GABA_*A*_ receptors exist as heteropentamers that contain two *α* and two *β* subunits, and either one *γ* or one *δ* subunit. Mainly due to this heterogeneity, a diverse set of ligands including anesthetics such as propofol and etomidate, and benzodiazepines such as diazepam, bind and modulate the functional characteristics of these channels [2–5].

Despite their biomedical relevance, and informative structures solved by cryoEM in apparent resting and desensitized states [6–11], structures of open *α*1*β*2*γ*2 GABA_*A*_ receptors have yet to be clearly defined. This lack is likely attributable to the transient time scale of the opening event (at the range of milliseconds [12–14]) which is much smaller than the sample preparation period of the cryo-EM grids. Even in the presence of different positive modulators such as anesthetics, benzodiazepines, and neurosteroids, a desensitized pore channel resulted as evidenced by the pore radius profiles that are smaller than that to permeate a hydrated chloride ion [6–11] (Fig. 1C, Fig. S1).

Molecular dynamics (MD) simulation offers complementary approaches to explore experimentally unseen states. However, conventional MD simulations fail to capture the timescale at which these channels function. Though enhanced sampling methods may address this problem [15], most of these methods require predefined knowledge of reliable reaction coordinates of the underlying biological process and the application of artificial biasing force. None of these criteria were satisfied in our case due to the 1) complex heteromeric assembly of the channel and 2) subtlety of the pore opening/closing which may differ as little as 1 Å between the open and desensitized states. To tackle these issues, a combination of adaptive sampling and Markov state modeling (MSM) has successfully been applied to exploring the protein conformational landscapes of systems such as Ebola viral protein 35 [16], epidermal growth factor receptor [17], and membrane transporters [18]. In particular, a goal-oriented adaptive sampling method named fluctuation amplification of specific traits (FAST) [19] was able to successfully predict an experimentally unseen open state of the SARS-CoV-2 spike complex [20]. It has yet to exploit the power of such methods to study ion channel structure and function.

Here, we applied FAST to first explore regions of conformational space relevant to the opening of the pore in the TMD of *α*1*β*2*γ*2 GABA_*A*_ receptor. MSM analysis of all the simulation trajectories revealed an energetically metastable state with a pore wide enough to conduct hydrated Cl^−^ ions and recapitulate previously measured experimental conductance value [21] (Fig. 1D,E). Furthermore, our analysis revealed the importance of the *αβ* interface in the activation of the channel; mutant simulations at this site indicated a decrease in the desensitization rate as seen in previous electrophysiology experiments (Fig. 1F) [14].

## Results

### Adaptive sampling widens the channel pore

To maximize the chance of discovering pore openings that allow ion permeation, we launched simulations from a cryoEM structure of the *α*1*β*2*γ*2 GABA_*A*_receptor determined in the presence of GABA and stabilizing Fab fragments (the latter removed in our simulation) in an apparent desensitized state (PDB ID: 6×3z [6]). We used an adaptive sampling method, FAST, that encourages the widening of the intersubunit distances at the TMD as the MD simulations progress [19] (See Methods for details) (Fig. 2A). We started simulations from the desensitized state, presuming it has a similar ECD conformation as in the open state, including the agonist binding. For efficiency, our simulations were designed to capture the transition between only open and desensitized states, with the resting state presumed to contribute relatively little in the presence of agonist. Within 30 FAST simulation generations, we observed around a 2-fold increase in hydration along the pore, associated with a widening of the pore (Fig. 2B).

**Fig. 2.**
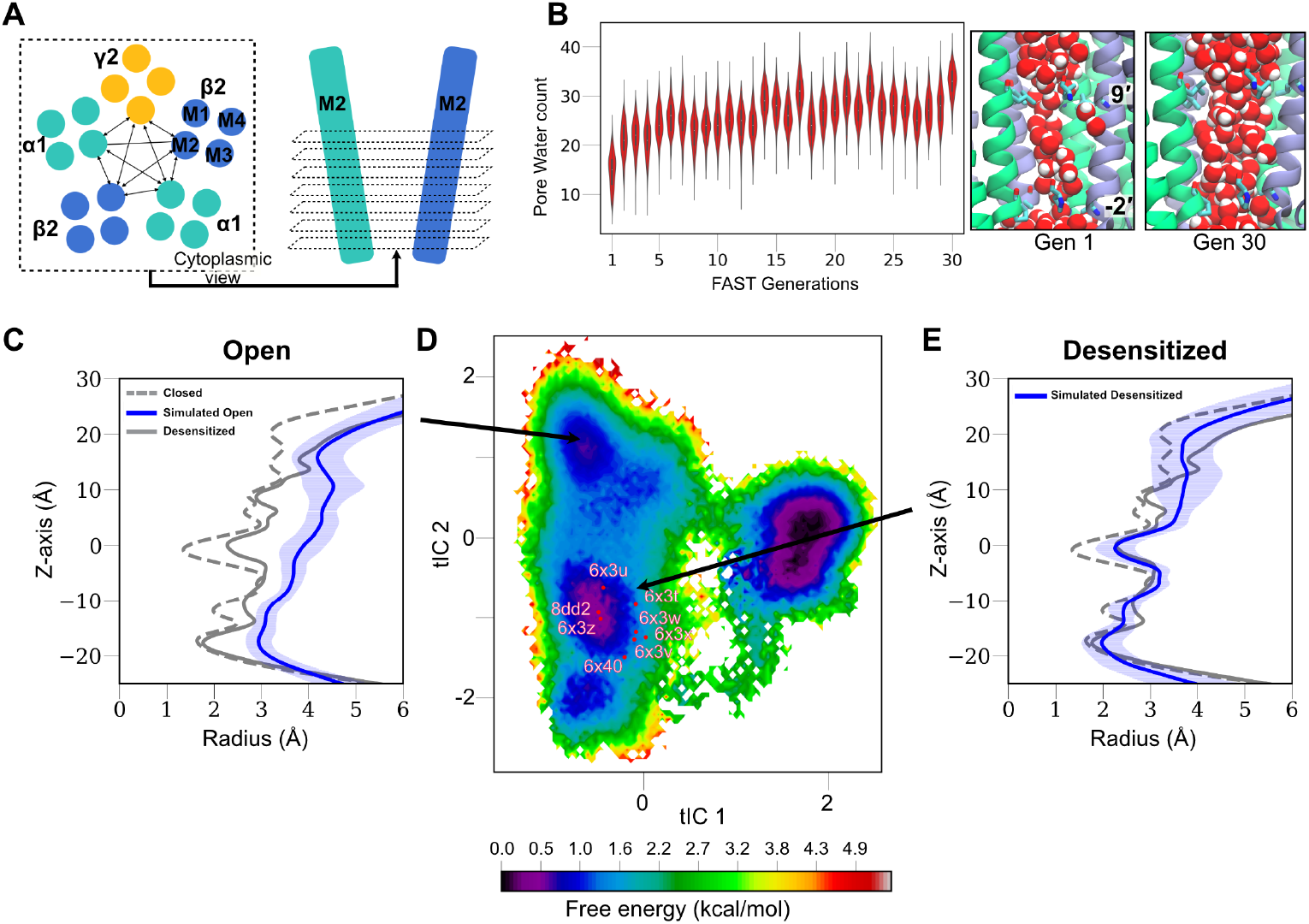
Structural and energetic characterization of the GABA_*A*_ receptor during our FAST sampling and MSM analysis. A) Schematic depiction of the feature selection criteria for the FAST sampling. All possible combinations of pair-wise distances among the M2 helix of all the subunits, spanning from the −4^*′*^ to 22^*′*^ were chosen as an input feature in FAST. B) Increased water hydration, along the progression of FAST generations, characterized by counting water in between the two gates (*left*). A representative snapshot of water hydration in the first and last generation of FAST sampling is shown on the right. C-E) Structural characteristics of the metastable states captured in the free energy landscape, projected onto the top two tICA eigenvectors from our MSM. The pore radius profile (calculated using CHAP [23]) with a mean (solid) and standard deviation (shaded) are shown in blue. The pore radius profile experimental closed (PDB ID: 6×3s) and desensitized (PDB ID: 6×3z) state structures are shown in dashed and solid gray, respectively. The structural feature of the third metastable state (rightmost) on the free energy landscape is shown in Fig. S4.

### Markov state modeling captures an energetically metastable open state

To identify key conformational states of GABA_*A*_ receptor, their underlying free-energy and kinetics of state transitions, we constructed an MSM using all the FAST-sampled MD trajectories (See Methods for details). The free energy was projected on the first two time-lagged independent components (tICs), which capture the slowest transitions of the system [24]. The energy landscape revealed 3 metastable states (Fig. 2C-E). Whereas all desensitized experimental structures projected to a well-associated desensitized state (described in detail below), our analysis revealed one putative open state in which both the central (9^′^) and intracellular (−2^′^) gates are wide enough for permeation of hydrated chloride ions, with an average radius of ≈ 4 Å and ≈ 3 Å respectively (Fig. 2C). Notably, MSM analysis showed that the open state is energetically less favorable, as expected relative to the long-lived desensitized state (Fig. 2C, D). This notion was further supported by our computed open state probability of 3.4%, using the first eigenvector of the transition probability matrix in our MSM analysis. A similarly low open state probability of around 16% was reported in previous experiments [25]. To further check the structural stability of this open state in additional MD simulations, we performed 6 replicates of simulations, each for 200 ns. In five of these replicates, the pore remained stably open, as evident by the minimal pore radius persisting at 3 Å at the end of the simulations (Fig. S2).

To test the dependence of our putative open state on the starting model, we repeated our FAST sampling followed by MSM analysis using the structure the *α*1*β*3*γ*2 GABA_*A*_ with alprazolam (PDB ID: 6huo). For consistency with our initial setup, all simulations were run with GABA alone as a ligand. The resulting free-energy landscape again contained a distinct well that corresponded to structures consistent with chloride conduction, and projecting these structures onto our initial landscape showed they largely corresponded to the same open state (Fig. S3).

The other two wells in our free energy landscapes were categorized as desensitized (Fig. 2E) and sheared (Fig. S4) states. In the desensitized well, the pore radius at both gates was similar to the experimental desensitized structure, with an average radius of ≈ 2 Å at the −2^′^ and ≈ 2.3 Å at the 9^′^ gate (Fig. 2E). In the sheared state, the 9^′^ gate was relatively expanded (3 Å radius), but the −2^′^ gate was again too narrow (2.2 Å radius) for a hydrated chloride to pass. Accordingly, we focused our subsequent analyses on the transition from predicted open to desensitized states.

### Simulated conductance of the open state agrees with experimental results

To validate the functional characteristics of our open-state model, we performed electric field simulations corresponding to an electric potential difference of 200 and -200 mV across the membrane, each for 6 replicates. These simulations revealed a conductance of 17 ± 8.9 pS at 200 mV and 9.3 ± 5 pS at -200 mV, similar to previous electrophysiology experiments reporting conductances values of 16.6 ± 3.6 (Fig. 3A) [21]. Note that at 200 mV, the chloride flows inward which is the predominant direction of ionic flow in neurons. Our desensitized model exhibited virtually no conductance at either voltage, confirming the presence of a constricted gate (Fig. 3A). We further characterized the energetics of chloride ion permeation from these two states by performing potential of mean force (PMF) calculations using the accelerated weight histogram (AWH) MD simulation method (Fig. 3B,Fig. S5) (details in the Method section) [26]. The ion faces lower energy barriers at both gates in the open state compared to the desensitized state, consistent with the higher conductance through the former model. Interestingly, in the open state, the highest barrier is still around 4 kcal/mol at the −2^′^ gate. This is possibly due to the partial dehydration, as defined by losing one water molecule in the first hydration shell, of the ion. This loss cannot be completely compensated by uncharged residues at this location (Fig. 3B, C).

**Fig. 3.**
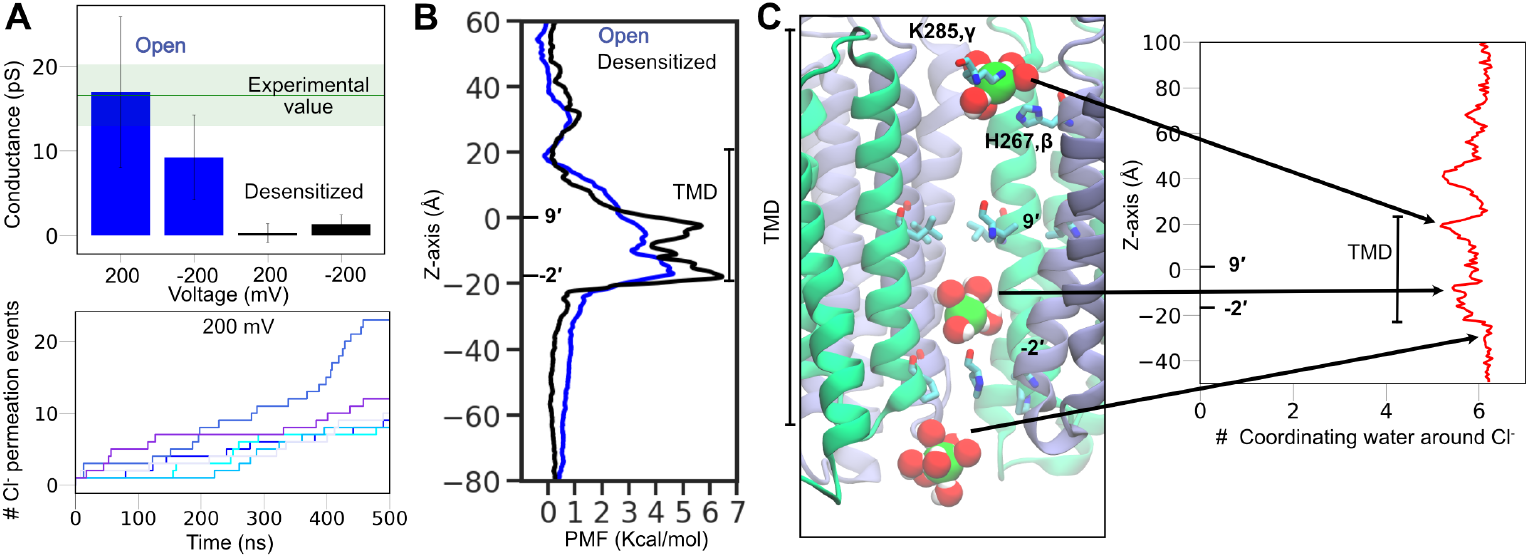
Simulated conductance of the open state agrees with experimental results. A, *top*) Mean and standard deviation of conductance for the open (blue) and desensitized (black) states, determined using six independent electric-field simulations for each state. The experimental values are depicted in a horizontal green line [21]. A, *bottom*) The cumulative net number of channel-crossing events by chloride ion, tracked over the time course of six independent MD simulations (colored differently) for the open state at 200 mV. B) Free energies for chloride ion permeation along the pore axis (with 9^*′*^ gate at 0 Å), for the open (blue) and desensitized state (black) of the receptor. The location of TMD is highlighted. The convergence of the free energies can be found in Fig. S5. C) Water coordination at the first hydration shell of chloride ion. Molecular snapshots at different locations of the ion are shown on the left. The water coordination number along the pore axis, calculated using all the simulation trajectories of the ion permeation free energy, is shown on the right.

### Open-to-desensitized transition is primarily mediated by the *αβ* interface

To extract mechanistic details of the open-to-desensitized transitions, we investigated subunit interactions at all five interfaces, focusing on residues around the −2^′^ and 9^′^ gates. At the −2^′^ gate, the desensitization process is primarily mediated by a pronounced contraction of the *αβ* interface, compared to the other interfaces (Fig. S6,Fig. 4A). A contraction of around 3 Å was observed for the *α*1*β*2*γ*2, with a similar trend for the *α*1*β*3*γ*2 system. This motion contracts a cleft at the interface between *α*-M2, M3 and *β*-M1, M2 helices, in the lower half of the TMD (Fig. 4B). Solvent accessible surface area (SASA) reduces around 20% at the *αβ* interface upon desensitization of the *α*1*β*2*γ*2 system, with a similar trend for *α*1*β*3*γ*2 (Fig. 4B). Notably, as captured in a previous cryo-EM structure (PDB ID: 6×3w), phenobarbital binds in the upper half of the TMD at this interface, a site that remains open in both states (Fig. S7). Around the 9^′^ gate, *α*-V263 (8^′^) at M2 holds away from *β*-V238 at M1 helix by around 2 Å during desensitization (with a similar movement for the *α*1*β*3*γ*2 system) and relaxes a kink at M2 (Fig. 4C, Fig. S8). This increase in the M2-M1 distance accompanied translocation of the 9^′^ residue towards the pore, narrowing the permeation pathway as described above (Fig. 2C).

**Fig. 4.**
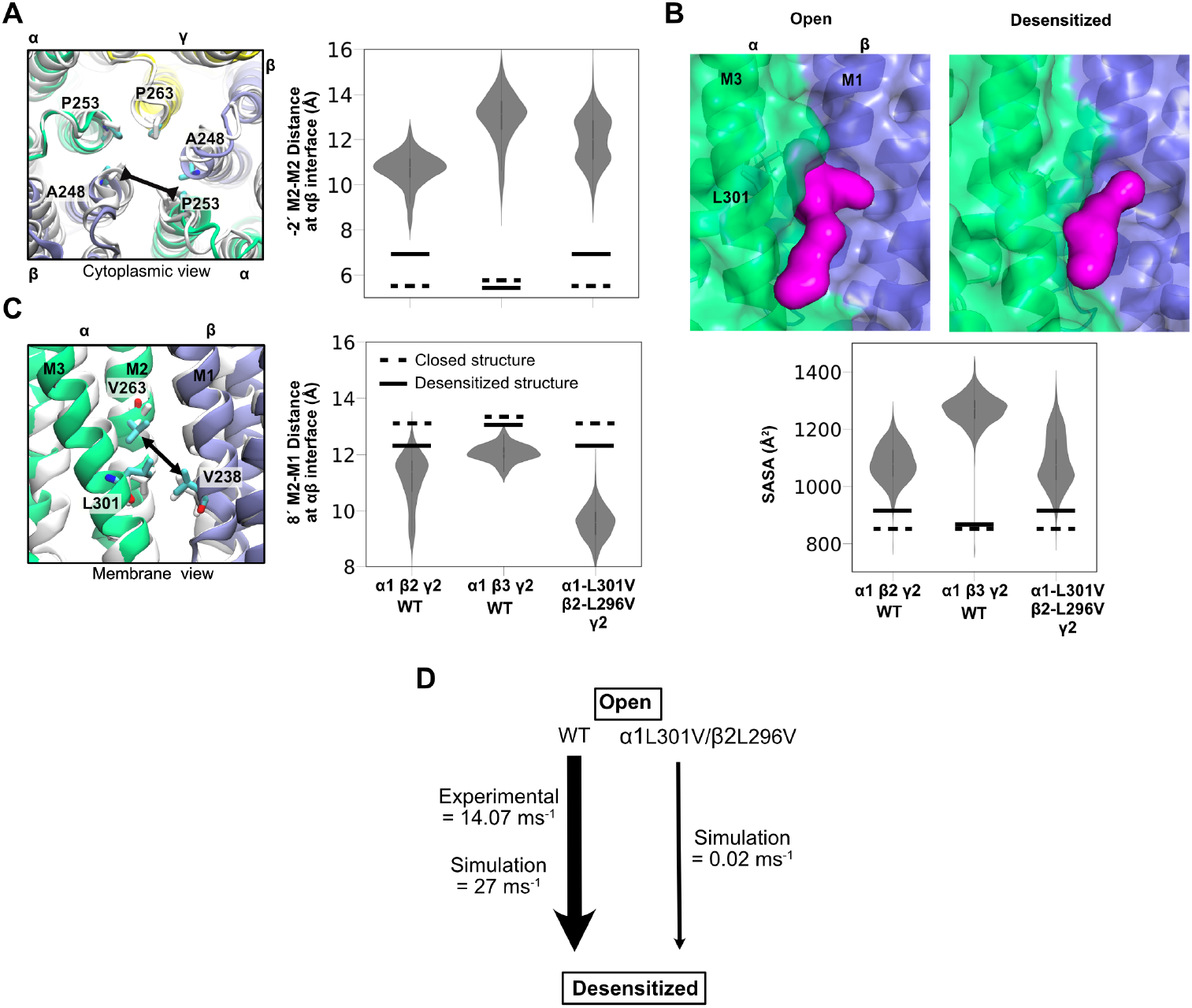
Open-to-desensitized transition mediated by the *αβ* interface and slowed by mutagenesis. A) Residue level details of the open state viewed from the cytoplasm at the −2^*′*^ gate, after aligning the entire TMD to the starting desensitized model (colored in white). Contraction at the *αβ* interface from our open states to the starting desensitized model is highlighted in the arrow. Distance between the C*α* atoms of the −2^*′*^ residues, located at the M2 helices of the interface, are depicted in the violin plot for the wild type (WT)-*α*1*β*2*γ*2, WT-*α*1*β*3*γ*2, and *α*1-L301V*β*2-L296V*γ*2 systems. The same distance matrix for the experimental closed and desensitized state structures are shown in the dotted and solid lines, respectively. B) Packing of the *αβ* interface calculated by measuring the solvent accessible surface area (SASA), revealing a possible opening of a druggable site (colored in purple) in the open state. Residue L301 (that is mutated in our study) is also highlighted. C) Molecular details of the *αβ* interface viewed from the membrane in the vicinity of the 9^*′*^ gate. The arrow represents the motion of the residue V263 (8^*′*^ location) residue, at the *α*-M2 helix, moving away from the residue V238, at the *β*-M1 helix, during the open-to-desensitization state transition. All distances were calculated using C*α* atoms. D) Open to desensitization transition rates, calculated for both the WT and the mutated system, using the transition path theory in pyEmma [27]. The line thickness qualitatively corresponds to the values of transition rates. Transition rates across all three states can be found in Fig. S9. The experimental value for the WT system is shwon [12].

### *In-silico* mutagenesis slows the open-to-desensitized transition, in accordance with experiments

Involvement of the M2–M1 intersubunit interface in desensitization echoed previous experimental studies. In particular, at the *αβ* interface, a valine substitution at a conserved leucine residue (*α*1-L301, *β*2-L296) located just past the midpoint of *α*-M3 has been shown to decrease desensitization rates in electrophysiology measurements (Fig. 2B,C) [14]. This M3 position faces *α*-V263 in M2 and *β*-V238 in M1, as well as the open-state crevice described above. To test if the *α*1-L301V/*β*2-L296V mutations can alter the desensitization process in our simulations, we introduced generated a new model by in-silico mutagenesis of the same GABA-bound desensitized channel GABA_*A*_receptor structure (PDB ID: 6×3z [6]) as used before. Then, we repeated our adaptive sampling method followed by MSM analysis.

The mutant open-state ensemble generally projected onto the wild-type free energy landscape (Fig. S3), and exhibited similar 1) asymmetric contraction along the *αβ* axis at the −2^′^ gate, 2) diminishing of a possible druggable pocket at the lower half of the TMD of the interface, and 3) expansion of the M2-M1 distance around the 9^′^ gate, while transiting from open-to-desensitized transition (Fig. 4A-C). Notably, the M2-M1 distance was even more contracted (by 2 Å) in the open state of the mutant compared to the wild-type (WT), possibly due to the space created by the L-to-V substitution (Fig. 4C).

To test if this pronounced M2-M1 interaction in the mutant open state can impact the desensitization process, we performed transition path analysis [28] using our MSM for both the WT and mutated system. In the WT system, we captured a desensitization timescale of 27 ms^−1^, comparable to the experimentally measured value of 14.07 ms^−1^ (Fig. 4D,Fig. S9) [12]. In the mutant system, a 1000-fold decrease in the open-to-desensitization transition rate was found, agreeing with the experimental findings (Fig. 4D) [14]. Strengthening of the M2-M1 interaction upon L301V substitution could discourage the 9^′^ residue from moving towards the pore, a process required for narrowing the permeation pathway and hence desensitizing the channel.

## Discussion

The open-state model generated here for the *α*1*β*2*γ*2 GABA_*A*_ receptor by an adaptive sampling/Markov-state modeling strategy is representative of a free-energy basin (Fig. 2C,D), metastable in extended simulations (Fig. S2), and largely reproducible from multiple starting models (Fig. S3). Although yet to be directly visualized by laboratory methods, our open-state model forms a fully hydrated pore and conducts chloride ions to a similar extent as in electrophysiology experiments [21] (Fig. 3A). Furthermore, interfacial mutations recapitulate a decrease in the desensitization rate shown in previous experiments. This model could thus be used for testing new mechanistic hypotheses and potentially even developing new pharmacological tools.

The channel activation and desensitization require both −2^′^ and 9^′^ gates to expand and contract, respectively (Fig. S10). A similar trend is seen among experimental structures of other members of the pentameric ligand-gated ion channels (pLGICs) family, namely *α* 1 glycine [29] and *α* 7 nicotinic acetylcholine receptors [30] (Fig. S10), suggesting a general gating mechanism of this family. This mechanism may represent a modification to the “dual gate” model, where activation can be thought to require expansion of only the 9^′^ gate, and desensitization to require contraction of only the −2^′^ gate [1].

We found two major motions at the *αβ* interface of the GABA_*A*_ receptor that regulate the channel gating (Figs. 4 and 5). At the −2^′^ gate, the interfacial M2 helices expand during activation and contract during desensitization. Around the 9^′^ gate, the M2(*α*)-M1(*β*) interface contracts during activation and expands during desensitization. A similar change in M2-M1 distance has been reported as an important feature in characterizing channel gating of various pLGICs [31]. The opposing motion at the two gates also introduces a slight kink at the 8^′^ residue. Interestingly, a similar M2 helix distortion was also captured recently in a PNU-bound open structure of the *α*7 nicotinic receptor [32].

**Fig. 5.**
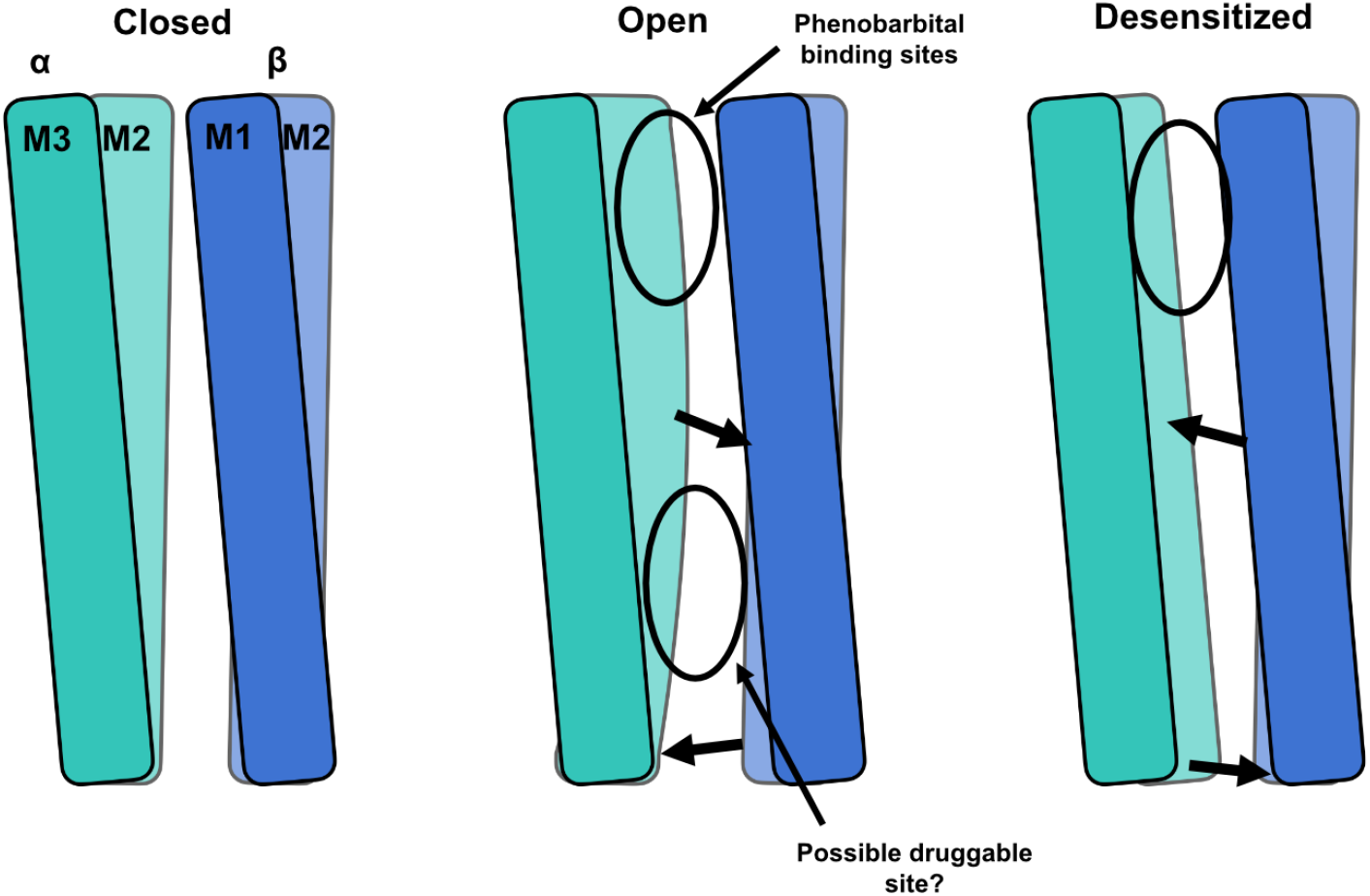
A structural model of *α*1*β*2*γ*2 GABA_*A*_ receptor gating proposed in this study.

Channel activation introduces two pockets at the *αβ* interface, formed by the M2 and M3 helices of the *α* and M1 and M2 helices of the *β* subunit (Figs. 4 and 5). The site at the upper half of the TMD has already been captured in cryo-EM with a positive modulator, phenobarbital-bound [6], although in a desensitized state (Fig. S7). To our knowledge, the site in the lower half of the TMD has yet to be implicated in binding any known modulators, suggesting a pathway for future development.

In addition to mechanistic details, our study provides thermodynamic and kinetic measures of channel function that can be compared to experiential findings. As seen in Fig. 2D, the free energy of the open state is relatively high, corresponding to an open probability of 3.4%, reminiscent of experimental values of 16% [25]. Our transition path analysis shows that the receptor desensitizes from the open state with a rate of around 27 ms^−1^, qualitatively agreeing with values from single-channel recordings of 14.07 ms^−1^ [12]. Although not perfectly precise, such measures offer benchmarks for further refinement of structural and mechanistic models.

We note the free energy basin for the sheared state is relatively lower than all the other states, which is surprising given the fact that none of the cryo-EM strictures belong to this metastable state. While this state might be present in cryo-EM particles that are not yet able to be extracted by class averaging, another possibility is insufficient sampling by the FAST method. Indeed, as expected we did not capture the closed state, judged by the fact that our MSM wells never sample a 9^′^ gate as narrow as in the experimental closed state (1.2 Å, Fig. 1C).

Overall, this study demonstrates the power of advanced sampling strategies and MSM analysis that can be used to address challenging biological questions. Given the fact that we did not use information on the open state from a homologous protein, this method appears to be a promising approach for sampling the conformational landscape of membrane protein in general, and in particular to propose testable models for novel functional states of ion channels.

## Methods

### System preparations

Simulations were initiated from the GABA-bound desensitized-state cryo-EM structure of GABA_*A*_ receptor (PDB ID: 6×3z), embedded and equilibrated for 20 *μ*s with coarse-grain models, as in our previous study [6]. Briefly, the model was coarse-grained through the representation of about four heavy atoms as a single bead, using Martini Bilayer Maker in CHARMM-GUI [33]. Then, the protein was embedded in a symmetric membrane containing 44.4% cholesterol, 22.2% 1-palmitoyl-2-oleoyl-sn-glycero-3-phosphocholine (POPC), 22.2% 1-palmitoyl-2-oleoyl-sn-glycero-3-phosphoethanolamine (POPE), 10% 1-palmitoyl-2-oleoyl-sn-glycero-3-phospho-L-serine (POPS) and 1.1% phosphatidylinositol 4,5-bisphosphate (PtdIns(4,5)P2), as previously shown to approximate the neuronal plasma membrane [34]. The equilibrated system was then backmapped to an all-atom system. For details, please see the reference [6]. The system was then solvated with TIP3P water [35] and neutralized with 0.15 M NaCl to generate systems containing 270,000 atoms, with dimensions of 140 × 140 × 160 Å^3^. A slightly different approach was taken for preparing the GABA+alprazolam-bound receptor (PDB ID: 6huo) [10] system (but, alprazolam removed) since we did not have any coarse-grain model equilibrated membrane system. To allow for better sampling of lipids (with the same composition as the other systems), 25 independent replicas were built by randomly configuring initial lipid placement around the protein using the Membrane mixer plugin in VMD [36, 37].

The systems were energy minimized and then relaxed in simulations at constant pressure (1 bar) and temperature (310K) for 30 ns, during which the position restraints on the protein and ligands were gradually released. The restraints were directly used as recommended by CHARMM-GUI. Then, production runs were performed with a mild flat-bottom restraint of 20 kJ mol^−1^ nm^−2^ between the atoms of GABA and residues at the binding sites, to prevent sudden release of GABA from the binding sites as seen in our previous simulation.

### Adaptive sampling

To explore possible pore openings at the TMD, we applied a goal-oriented adaptive sampling method, called fluctuation amplification of specific traits (FAST) [19]. Briefly, the method runs successive swarms of simulations where the starting points for each swarm are chosen from the set of all previously discovered conformations based on a reward function. This function balances (1) preferentially simulating structures with maximum pair-wise distances (with a total of 540 pairs, Fig. 2A) to encourage the pore to adopt a more open conformation that may harbor ion permeation; with (2) a broad exploration of the conformational space of the entire receptor. The broad exploration phase was implemented by favoring states that are poorly sampled compared to other states, based on the root root-mean-square deviation of the TMD residues. During FAST, we performed 30 generations of simulations with 25 simulations/generation and 40 ns/simulation, totaling to 30 *μ*s. Since no biasing force is applied to any individual simulation, the final data set can be used to build a Markov state model (MSM) to extract the proper thermodynamics and kinetics [38–40], as detailed below.

### Markov state modeling

We used our trajectory dataset from FAST to construct a Markov state model (MSM) using pyEmma [27], by first featurizing the trajectory dataset using: 1) the pore radii, spanning from −2^′^ to 9^′^, every 1 Å, and 2) M2-M2 intersubunit distances and M2-M1 and M2-M3 intrasubunit distances spanning the same region as above. To remove redundant information within the feature space and identify the slowest reaction coordinates, time-structure based independent component analysis (tICA) was used to reduce the dimensionality of the feature space (*X*(*t*)) to the eigenvectors of an autocovariance matrix, ⟨*X*(*t*)*X*^*T*^ (*t* + *τ*)⟩, with a lag time, *τ* =1 ns [24, 41, 42]. It is important to choose an optimal number of tICA eigenvectors since an MSM built using too many eigenvectors would have microstates with low statistical significance due to finite sampling error [43]. We found that the first twelve tICA eigenvectors are sufficient to construct the MSM, as assessed by the convergence of VAMP-2 score (Fig. S11) [44]. The conformational space was then discretized into multiple microstates using k-means clustering. To choose the number of microstates to use in the model, we used the VAMP-2 score [44], to evaluate the quality of MSMs built with different numbers of microstates (Fig. S11).

Then, a transition probability matrix (TPM) was constructed by evaluating the probability of transitioning between each microstate within a lag time, *τ*. To choose an adequate lag time to construct a TPM that ensures Markovian behavior, multiple TPMs were first created using multiple maximum-likelihood MSMs with different lag times. The implied timescales were evaluated for each of these transition matrices, and saturation was observed at *τ* = 5 ns for the WT and *τ* = 8 ns for the mutant system (Fig. S12). Thus, we built our final TPM using a maximum likelihood MSM with the corresponding lag times. This final TPM is symmetrized using a maximum likelihood approach to ensure detailed balance [27].

### MD simulations

MD simulations in this study were performed using GROMACS-2023 [45] utilizing CHARMM36m [46] CHARMM36 [47] force field parameters for proteins and lipids, respectively. The force field parameters for the ligands were generated using the CHARMM General Force Field (CGenFF) [48–50]. Cation-*π* interaction-specific NBFIX parameters were used to maintain appropriate ligand-protein interactions at the aromatic cage, located at the binding sites [51]. Bonded and short-range non-bonded interactions were calculated every 2 fs, and periodic boundary conditions were employed in all three dimensions. The particle mesh Ewald (PME) method [52] was used to calculate long-range electrostatic interactions with a grid density of 0.1 nm^−3^. A force-based smoothing function was employed for pairwise nonbonded interactions at 1 nm with a cutoff of 1.2 nm. Pairs of atoms whose interactions were evaluated were searched and updated every 20 steps. A cutoff of 1.2 nm was applied to search for the interacting atom pairs. Constant pressure was maintained at 1 bar using the Parrinello-Rahman algorithm [53]. Temperature coupling was kept at 300K with the v-rescale algorithm [54].

### Ion permeation free energy calculations

The free energy along the pore axis for chloride was calculated using the accelerated weight histogram (AWH) method [26]. In brief, for each structural model of open and desensitized state captured in our MSM analysis, we applied one independent AWH bias and simulated for 500 ns each with 8 walkers sharing bias data and contributing to the same target distribution. Each bias acts on the centre-of-mass z-distance between one central chloride ion and the −2^′^ residues, with a sampling interval across more than 95% of the box length along the z-axis to reach periodicity. To keep the solute close to the pore entrance, the coordinate radial distance was restrained to stay below 10 Å by adding a flat-bottom umbrella potential. During these simulations, the protein-heavy atoms were harmonically restrained using a relatively weak force constant of 100 kjoule mol^−1^ Å^−2^ to the backbone atoms to maintain the respective conformational state of the protein.

### Electric field simulations

Ionic current was calculated by performing simulations with a constant electric field normal to the membrane. Six replicas of ionic current simulations were performed with an open and desensitized state GABA_*A*_ receptor model, derived from MSM analysis. Each production simulation was then performed for 500 ns with an electric field corresponding to a membrane electric potential difference of 200 and -200 mV. During these simulations, the protein-heavy atoms were harmonically restrained using a relatively weak force constant of 100 kjoule mol^−1^ Å^−2^ to the backbone and 20 kjoule mol^−1^ Å^−2^ to the side-chain atoms to maintain the respective conformational state of the protein.

Ionic current (I) was computed by counting the number of ions (Na^+^ and Cl^-^) that cross the porin over time, i.e., 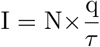, where N is the number of ion crossing events over a time interval *τ*, and q is the charge of the ion (1.60217662×10^−19^ Coulombs for Na^+^, and -1.60217662×10^−19^ Coulombs for Cl^-^). The total current was simply the sum of the net Na^+^ current minus the net Cl^-^ current. The conductance (C) was then calculated as 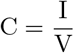.

### Analysis

System visualization and analysis were carried out either using VMD [37] or PyMOL [55].

## Acknowledgments

We thank Maxwell Zimmerman, Ryan Hibbs, and Weronika Chojnacka for their valuable feedback and discussion. MD simulations were performed using computing facilities of the Karolina, LUMI and Discoverer Supercomputer through EuroHPC (grant nos. EHPC-REG-2023R01-103, EHPC-REG-2022R03-223 and EHPC-REG-2022R03-219, respectively) and the Swedish National Infrastructure for Computing (SNIC 2022/3-40) and supported by BioExcel (EuroHPC grant no. 101093290). N.H. was supported by a Marie Sklodowska-Curie Postdoctoral Fellowship (grant no. 101107036), R.J.H., and E.L. by grants from the Swedish Research Council (2019-02433, 2021-05806) and Swedish e-Science Research Center.

## Data and code availability

The raw MD simulation trajectories can be found on Zenodo doi 10.5281/zenodo.10964268.

## Author contributions

N.H. designed research, performed the simulations and analyzed data; and N.H., R.J.H., and E.L. wrote the paper. All authors read and contributed to finalizing the paper.

