## Supplementary Information for "Adaptive sampling-based structural prediction reveals opening of a GABA_*A*_ receptor through the *αβ* interface"

### Supplementary Figures

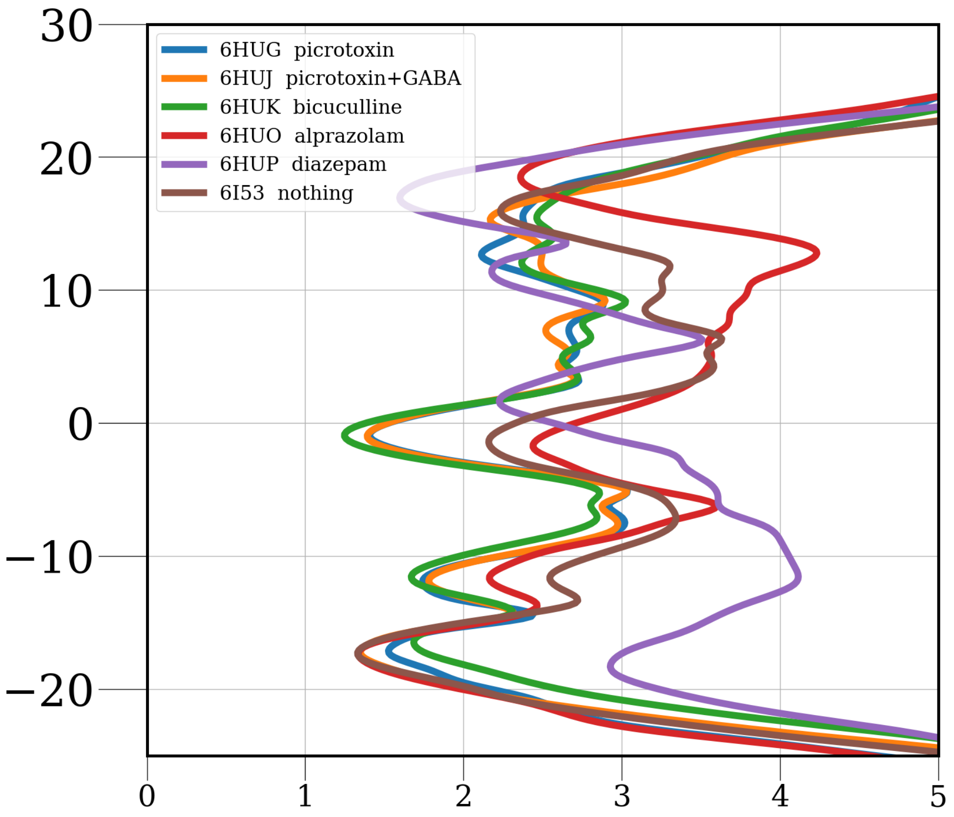


**Fig. S1** Pore radius profile of $\alpha 1\beta 3\gamma 2$ GABA$_A$ receptor structures, either in the closed or desensitized state, bound to different ligands [10, 11].

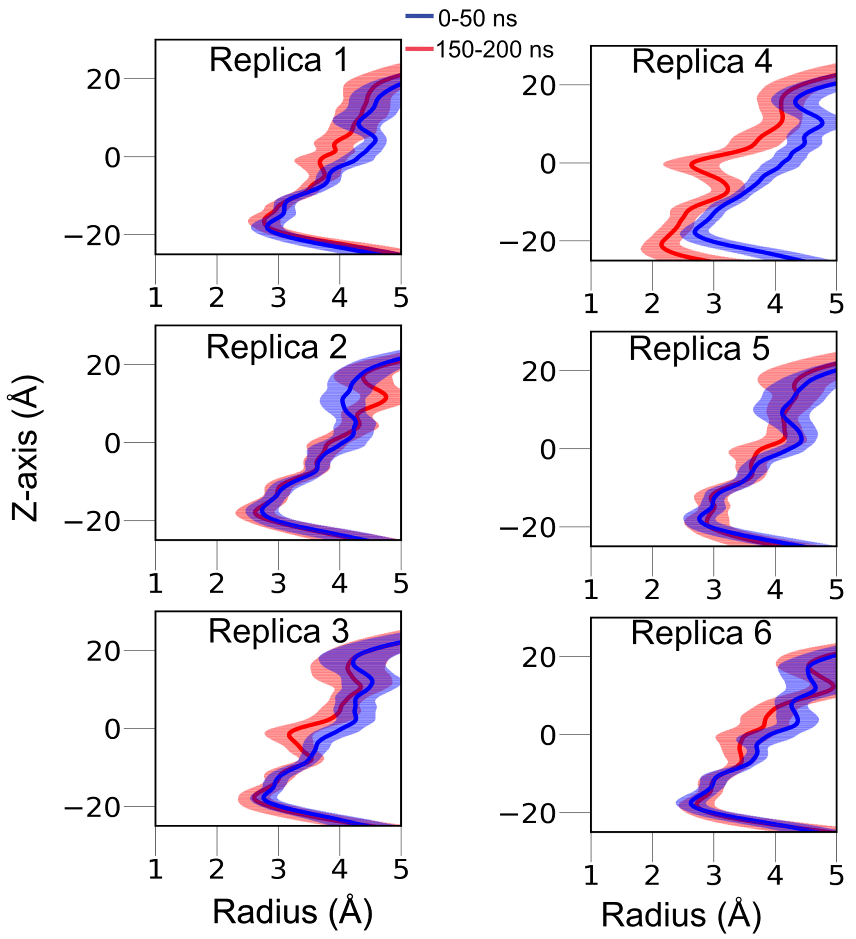


**Fig. S2** The pore radius profile with a mean (solid) and standard deviation (shaded) are shown for the first (blue) and last (red) 50 ns, for all the six replicates of MD simulations, starting from a representative open state model from our MSM (Fig. 2). In five out of six replicates, the pore remains stably open.

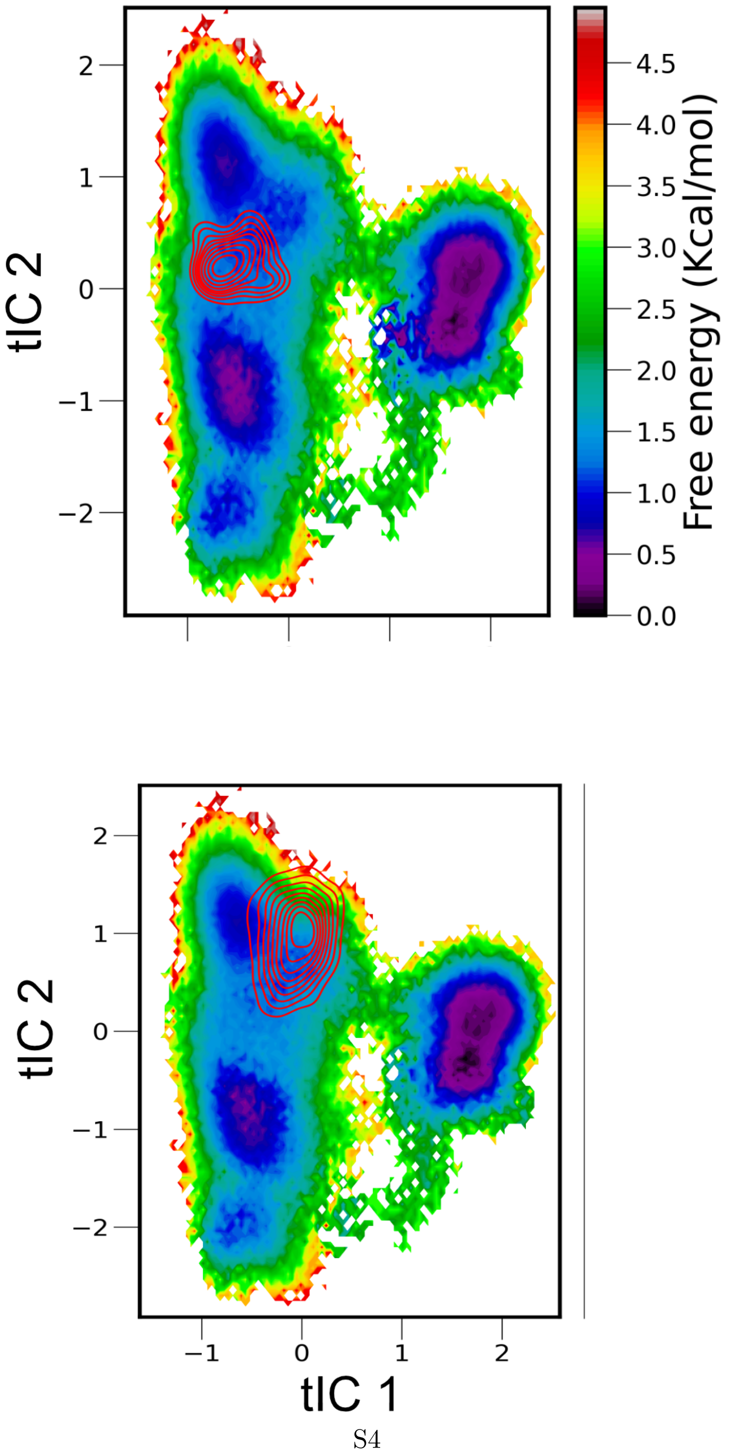




**Fig. S3** Projection of open state models captured in WT $\alpha1\beta3\gamma2$ (*top*) and mutant $\alpha1$-L301V$\beta2$-L296V$\gamma2$ (*bottom*) systems using FAST sampling followed MSM analysis, on the MSM of the WT $\alpha1\beta2\gamma2$ GABA$_A$.

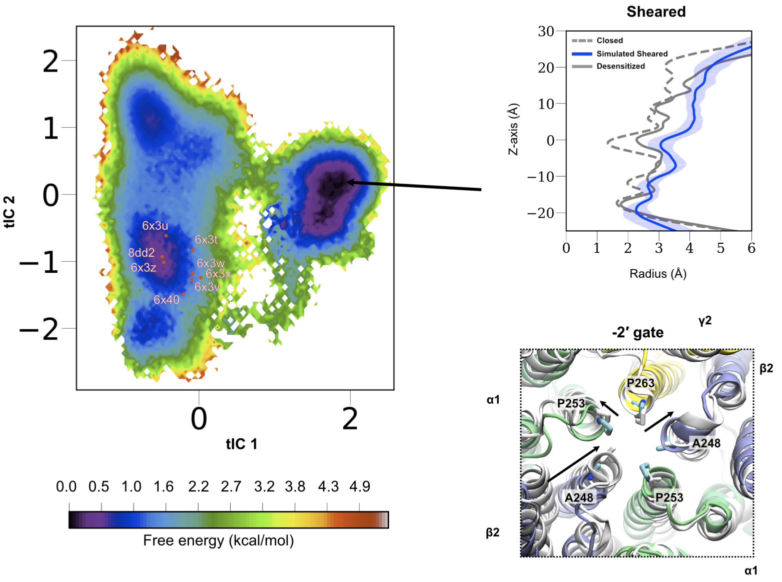


**Fig. S4** Structural characteristics of the metastable rightmost sheared state captured in the free energy landscape, projected onto the top two tICA eigenvectors from our MSM. The pore radius profile (calculated using CHAP [23]) with a mean (solid) and standard deviation (shaded) are shown in blue. The pore radius profile experimental closed (PDB ID: 6x3s) and desensitized (PDB ID: 6x3z) state structures are shown in dashed and solid gray, respectively. One of the $\beta$ subunits appears "displaced" by around 1 Å (as measured by the $\beta$-$\beta$ distance at the $-2'$ gate) from the pore, while the other $\alpha$ subunit by 0.5 Å (as measured by the $\alpha$-$\alpha$ distance) and a nearby $\beta$ subunit (as measured by the $\alpha$-$\beta$ distance, across the pore) collapsed towards the pore.

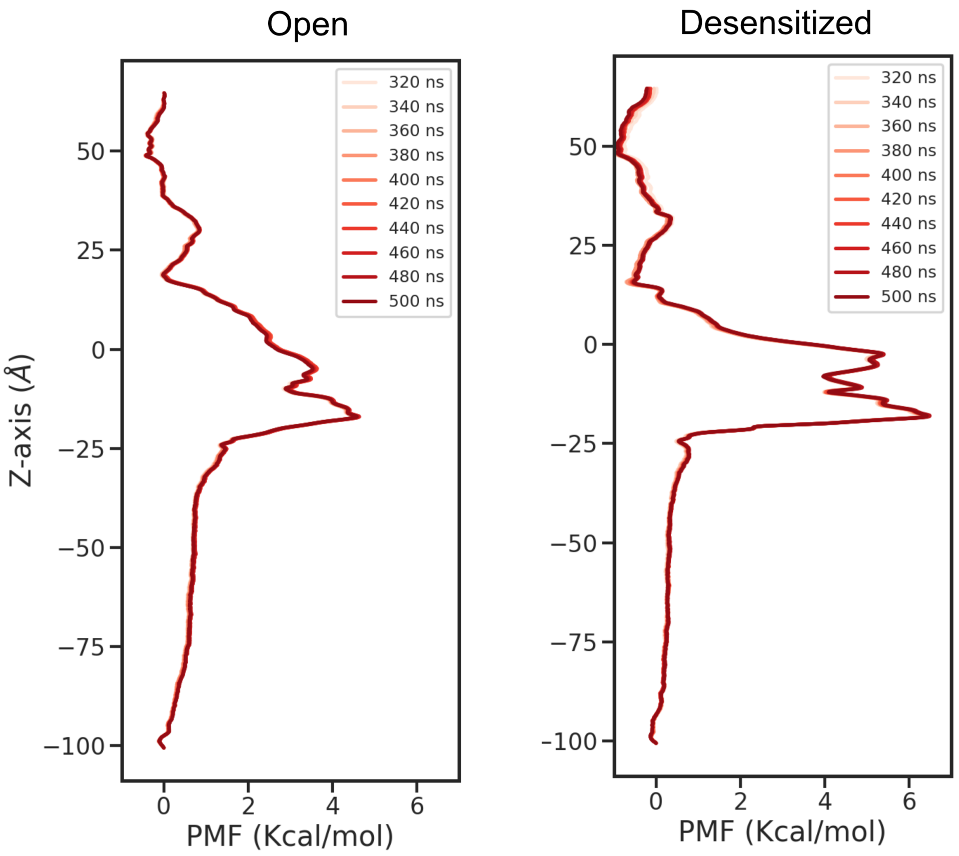


**Fig. S5** Convergence of ion permeation free energy calculations for both open and desensitized states.

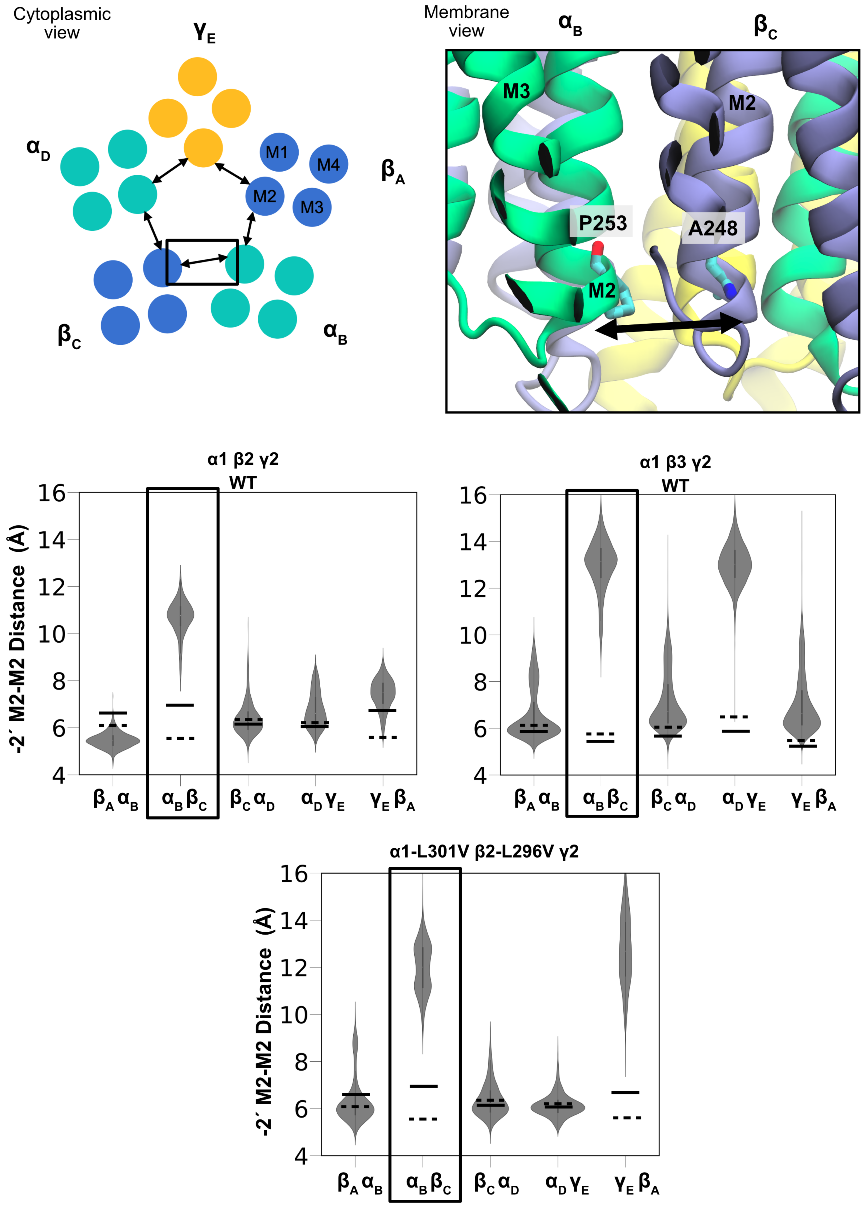


**Fig. S6** Intersubunit distances at the $-2'$ gate, calculated by selecting the C$\alpha$ atoms of the corresponding residues at this location at each subunit. Schematic depictions for each distance vector are shown above. The violin plot of these distance distributions for the open state captured for each of the three systems investigated in this study is shown. The corresponding distances for the experimental closed and desensitized state structures are shown in the dotted and solid lines, respectively.

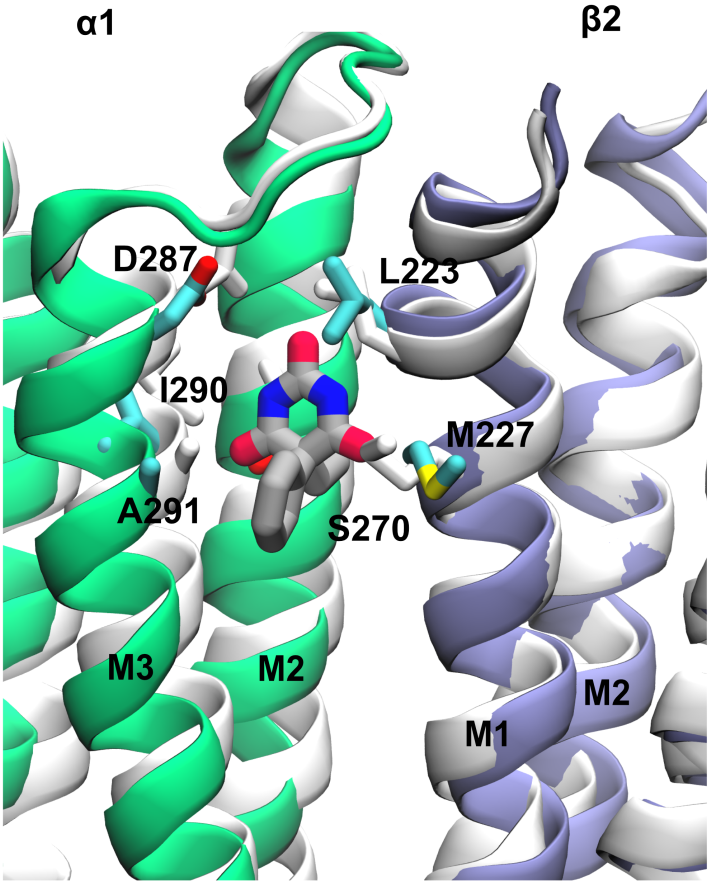


**Fig. S7** Phenobarbital drug site, resolved in cryo-EM structure of the $\alpha1\beta3\gamma2$ GABA$_A$ (PDB ID: 6X3W). The modulator-unbound experimental desensitized and simulated open states are shown in white and color, respectively.

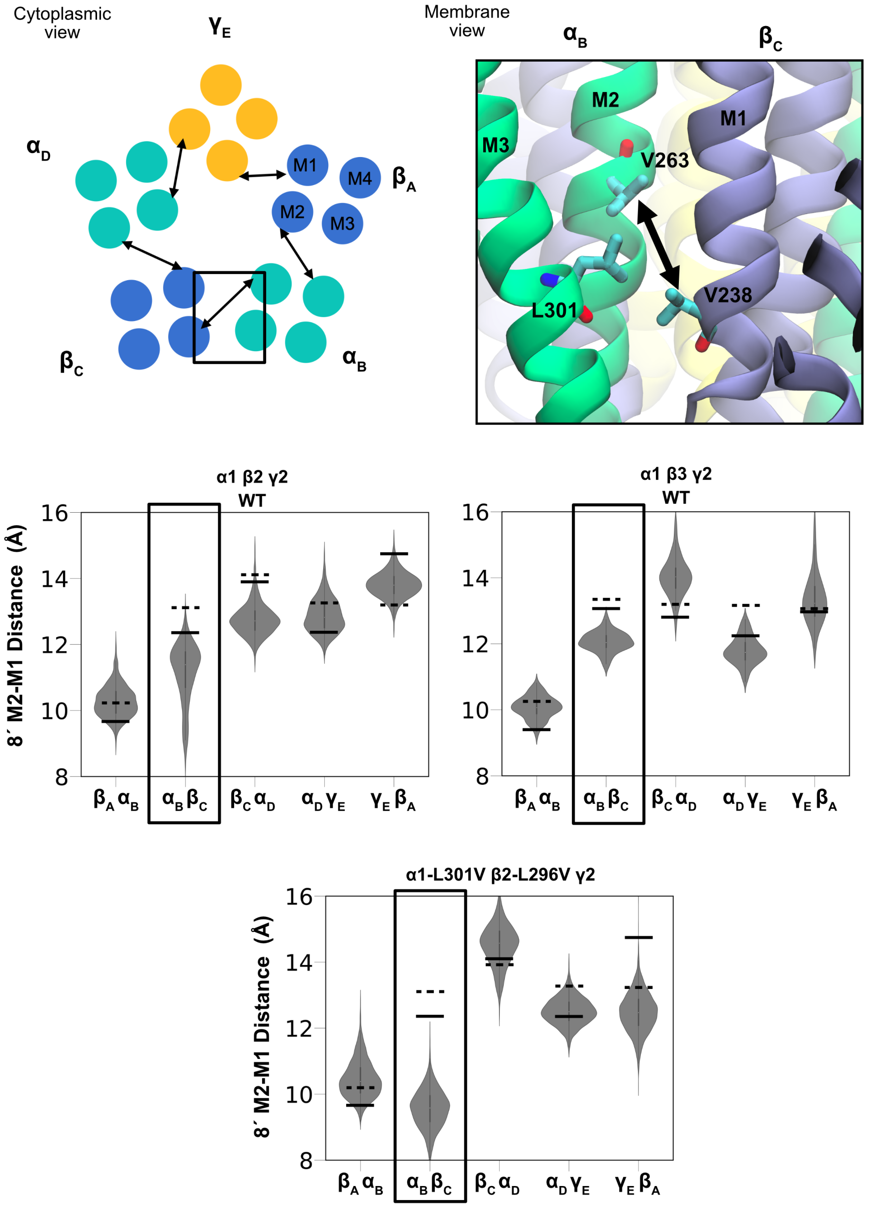


**Fig. S8** Intersubunit M2-M1 distances nearby the 9′ gate. The distances were calculated between the residue V263 (8′ location), at the $\alpha$-M2 helix, and V238, $\beta$-M1 helix. Schematic depictions for each distance vector are shown above. The violin plot of these distance distributions for the open state captured for each of the three systems is shown. The corresponding distances for the experimental closed and desensitized state structures are shown in the dotted and solid lines, respectively.

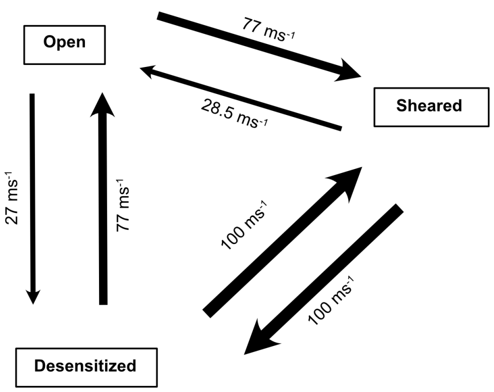


**Fig. S9** Transition rates, calculated for in and out of each state of the WT system.

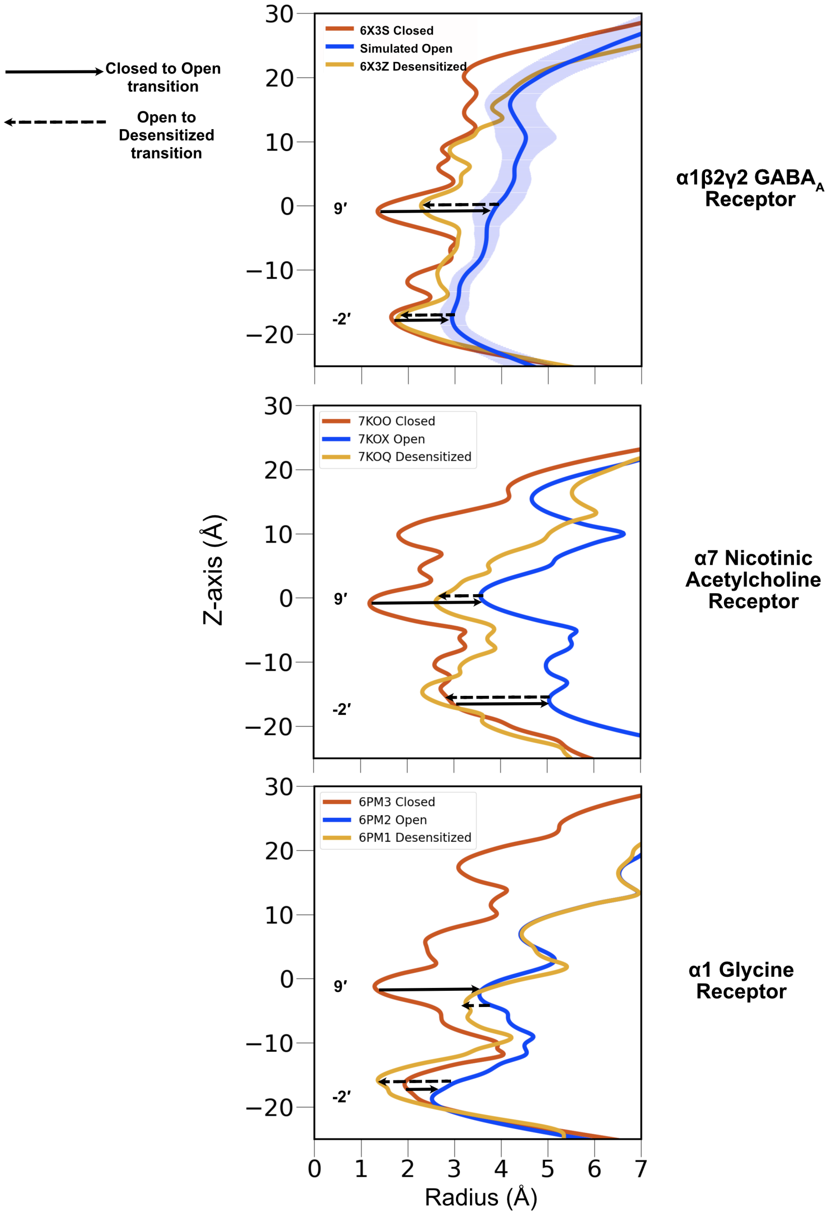


**Fig. S10** Pore radius profile of three different pLGIC receptor structures, in the closed, open and desensitized state. For all three receptors, expansion of the $-2'$ and $9'$, transiting from the closed to open state, and contraction of both gates transiting from open to desensitized state are depicted by solid and dashed arrows, respectively.

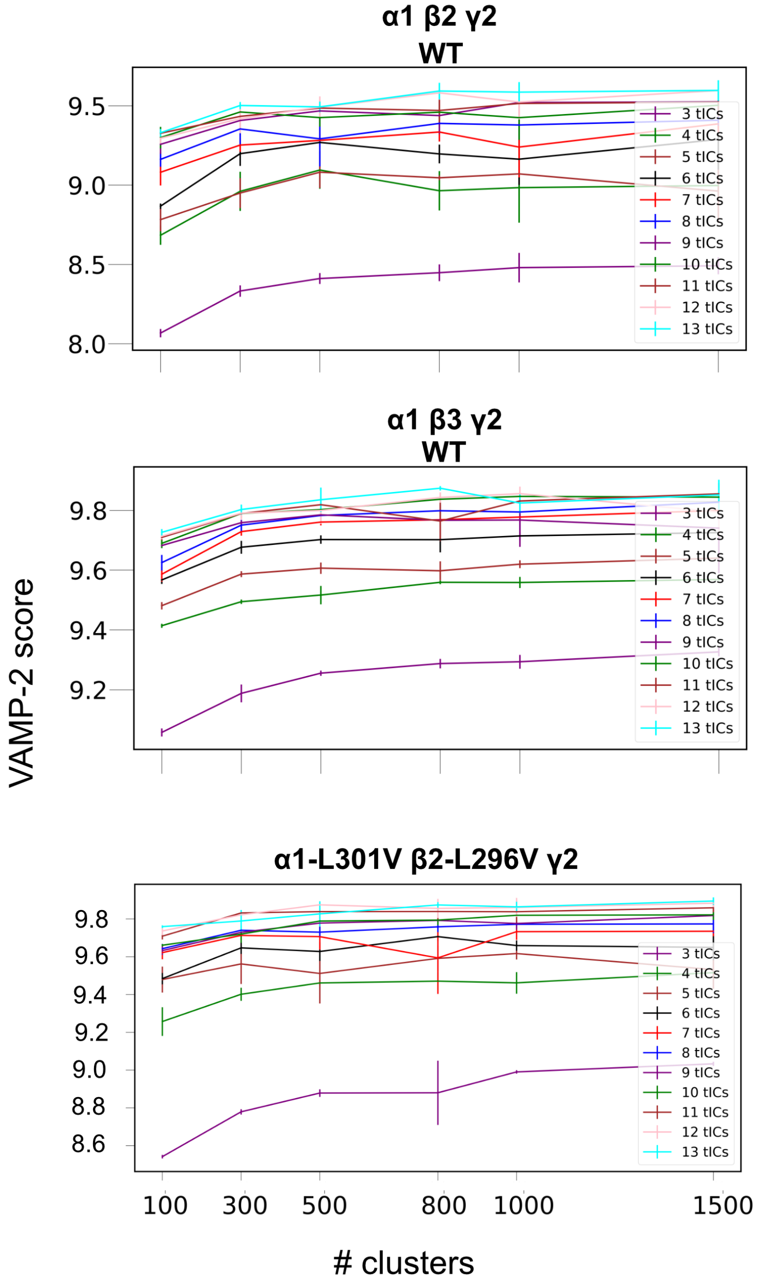


**Fig. S11** Cross-validated VAMP-2 [44] score to rank the parameters (number of tICA eigenvectors and clusters) used to discretize the conformational space. Calculations are performed independently for each system. Error bars are calculated by determining the VAMP-2 score after running 5 iterations of the k-means clustering algorithm.

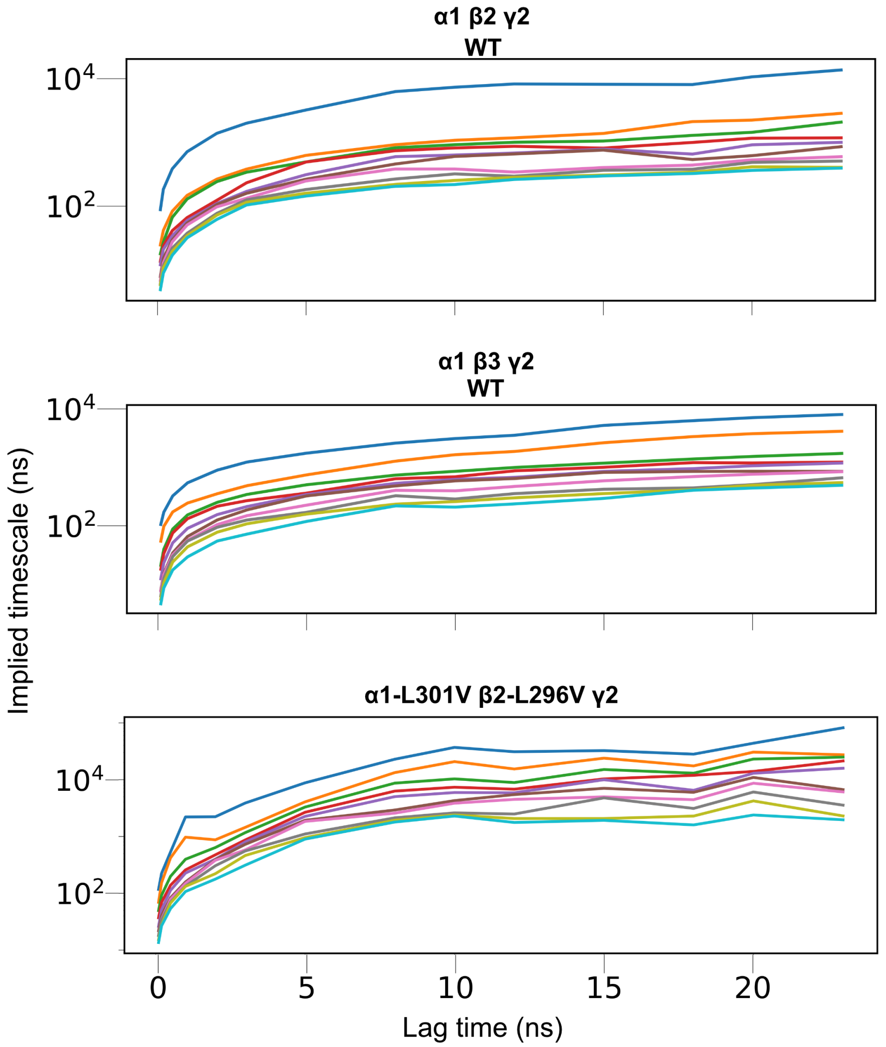


**Fig. S12** Implied timescales (ITS) plot of the top 10 slowest processes using multiple lag times, calculated independently for each system. The lag time of 5 ns and 8 ns were chosen for MSM construction for WT and mutant systems, respectively, as the ITS curve converges, indicating Markovianity.
